# The guanine nucleotide exchange factor GBF1 participates in rotavirus replication

**DOI:** 10.1101/619924

**Authors:** José L. Martínez, Francesca Arnoldi, Elisabeth M. Schraner, Catherine Eichwald, Daniela Silva-Ayala, Eunjoo Lee, Elizabeth Sztul, Óscar R. Burrone, Susana López, Carlos F. Arias

## Abstract

Cellular and viral factors participate in the replication cycle of rotavirus. We report that the guanine nucleotide exchange factor GBF1, which activates the small GTPase Arf1 to induce COPI transport processes, is required for rotavirus replication since knocking down GBF1 expression by RNA interference, or inhibiting its activity by treatment with Brefeldin A (BFA) or Golgicide A (GCA) significantly reduce the yield of infectious viral progeny. This reduction in virus yield was related to a block in virus assembly since in the presence of either BFA or GCA the assembly of infectious mature triple-layered virions was significantly prevented and only doubled layered-particles were detected. We report that the catalytic activity of GBF1, but not the activation of Arf1, is essential for the assembly of the outer capsid of rotavirus. We show that both BFA and GCA, as well as interfering with the synthesis of GBF1, alter the electrophoretic mobility of glycoproteins VP7 and NSP4 and block the trimerization of the virus surface VP7, a step required for its incorporation into virus particles. Although a post-translational modification of VP7 (other than glycosylation) could be related to the lack of trimerization, we found that NSP4 might also be involved in this process, since knocking-down its expression reduces VP7 trimerizarion. In support, recombinant VP7 protein overexpressed in transfected cells formed trimers only when co-transfected with NSP4.

**IMPORTANCE:** Rotavirus, a member of the family Reoviridae, is the major cause of severe diarrhea in children and young animals worldwide. Despite the significant advances in the characterization of the biology of this virus, the mechanisms involved in morphogenesis of the virus particle are still poorly understood. In this work, we show that the guanine nucleotide exchange factor GBF1, relevant for the COPI/Arf1-mediated cellular vesicular transport, participates in the replication cycle of the virus, influencing the correct processing of viral glycoproteins VP7 and NSP4, and the assembly of the virus surface proteins VP7 and VP4.

## INTRODUCTION

Rotaviruses, members of the family *Reoviridae*, are non-enveloped particles formed by three concentric layers of proteins that surround the eleven genome segments of double-stranded RNA (dsRNA). The innermost layer is composed of the core-shell proteinVP2 that encloses the replication intermediates, composed of the RNA dependent RNA polymerase VP1, and the guanylyl-methyl transferase, VP3. The intermediate layer is formed by VP6 that surrounds the VP2 layer to form double-layered particles (DLPs). Finally, the addition of the glycoprotein VP7 and the spike protein VP4 onto the DLPs forms the infectious triple-layered particles (TLPs) (1, 2).

The replication of rotavirus occurs in cytoplasmic non-membranous electron-dense inclusions termed viroplasms composed of NSP2, NSP5, VP1, VP2, VP6 and host components (1, 3). The replication and packaging of the viral genome into newly synthesized DLPs take place in these inclusions (4), which then bud into the lumen of the endoplasmic reticulum (ER) through membrane sites modified by the presence of NSP4 (5, 6). NSP4 is a transmembrane ER glycoprotein with two N-linked high mannose glycosylated chains (7) that play a crucial role in the last steps of rotavirus assembly. It has been shown that the cytoplasm oriented-terminus of NSP4 associates with VP4 (8), and binds the VP6 on DLPs acting as a receptor for these particles to mediate their budding into the ER (9, 10). Moreover, NSP4 has been shown to also interact with VP7 through its N-terminus oriented to the ER lumen (11, 12). It has been proposed that these interactions drive the incorporation of the outer layer proteins into the transitory lipid envelope that DLPs acquire during ER membrane budding; this envelope is removed in the lumen of the ER by an unknown process in which NSP4 is eliminated while VP4 and VP7 are assembled to produce the final infectious TLPs (13, 14). Although the precise mechanism of the final steps of rotavirus assembly is not well understood, it has been found that VP7 structure forms trimers on the surface of the virion in a calcium-dependent process (15–18).

Due to the high complexity of rotavirus replication, many of the cellular factors and molecular mechanisms involved in this process are poorly characterized. However, it has been recently reported that the coatomer protein I (COPI)/Arf1 machinery is essential for virus replication since knocking-down by RNA interference (RNAi) the expression of some of the proteins that integrate such machinery reduces virus replication (19, 20). Also, brefeldin A (BFA), an inhibitor of the COPI/Arf1-mediated vesicular transport, significantly impairs the rotavirus progeny yield (21). COPI is a protein complex formed by seven subunits (α, β, β’, δ, ε, γ and ζ-COP) that mediates the retrograde transport of vesicles from the Golgi apparatus to the ER (22–24). Besides its canonical functions in the secretory pathway, the COPI/Arf1 machinery also may participate in the maturation of early endosomes and in recycling proteins to the plasma membrane (25, 26), as well as in the maturation of phagosomes (27, 28) and peroxisomes (29). Furthermore, multiple reports suggest that the COPI/Arf1 machinery is also involved in transport events involved in the maturation and function of lipid droplets (LDs) (30–32).

In the initial step of the COPI transport, the small GTPase Arf1 (ADP-ribosylation factor 1) is activated with a GTP molecule in a process catalyzed by the guanine nucleotide exchange factor GBF1 (Golgi-specific BFA resistance factor 1) located at the cis-Golgi membrane and the intermediate ER-Golgi compartment (ERGIC) (33). The activated Arf1-GTP associates with the Golgi membrane and promotes the recruitment of the preformed COPI complex as well as of the Arf1-GTPase-activating protein (Arf1GAP). The formation of Arf1-COPI-Arf1GAP complex stimulates the binding and concentration of different cargoes located in the membrane, association that induces the bending of the membrane into a vesicle. Once completed, the vesicle buds from the membrane covered by the COPI complex proteins. Finally, the coat proteins are disassembled when the GTPase activity of Arf1 is enhanced by Arf1GAP, leading to the hydrolysis of the GTP bound to Arf1. This hydrolysis leads to the release from the membrane of Arf1, COPI, and Arf1GAP (34, 35).

GBF1 belongs to a subfamily of large guanine nucleotide exchange factors (GEFs) that also includes the mammalian BIG1, and BIG2 located at the trans-Golgi network (TGN) (36). These three GEFs activate Arf1, however GBF1 may also use Arf4 and Arf5 as substrates (37–39), while BIG1 and BIG2 can catalyze the activation of Arf3, Arf5 and Arf6 (40–42). Arf activation is catalyzed by the Sec7 domain, shared by all GEFs. In addition, GBF1 contains five non-catalytic domains: the N-terminal dimerization and cyclophilin binding (DCB), the homology upstream of Sec7 (HUS), and three C-terminal downstream of Sec7 (HDS1-3) domains (33, 36). The functions of these non-catalytic domains are not well understood, but the N-terminal DCB and HUS domains have been implicated in inter- and intra-molecular interactions important for GBF1 dimerization and its association to membranes (43, 44), while the HDS1-3 domains have been suggested to facilitate GBF1 association with membranes (Bouvet et al., 2013; Chen et al., 2016; Ellong et al., 2011; Meissner et al., 2018 and unpublished results).

In this study, we evaluated the role of the GBF1/COPI/Arf1 machinery in rotavirus replication. We showed that the catalytic activity of GBF1 is critical for virus replication by using two different pharmacological inhibitors, BFA and Golgicide A (GCA), and knocking down GBF1 expression by RNAi. We found that interfering with GBF1 activity significantly impaired the yield of viral progeny through a block in the assembly of the virus surface proteins, VP7 and VP4, which prevents the production of mature and infectious TLPs. We also showed that this restriction in the assembly of TLPs is the result of a damage in VP7 trimerization required for its assembly into DLPs. We provide evidence suggesting that an altered post-translational modification (different from glycosylation) of either VP7 or NSP4 in GBF1-inactivated cells is responsible for the defective formation of VP7 trimers. Altogether, our findings suggest that GBF1 activity is essential for the rotavirus outer capsid assembly by allowing the correct processing of VP7 and/or NSP4, possibly through a mechanism independent of Arf1.

## RESULTS

### Inhibition of GBF1 hinders rotavirus replication

Earlier reports have indicated that the COPI/Arf1 machinery is important for rotavirus replication (19, 20) and we characterized the effects of the pharmacological inhibitors BFA and GCA, which block the GBF1-mediated activation of Arf1 required for COPI transport (34, 35, 49, 50) on the replication of rotaviruses. In these assays, MA104 cells were pretreated for 30 min with different concentrations of BFA or GCA before virus infection. The simian rhesus rotavirus (RRV) was then added to cells in the presence of the inhibitors for 1 h at 37°C. Then, the virus inoculum was removed, and fresh media containing the inhibitors were added; 12 h post infection (hpi), cells were harvested, and the viral yield was determined. We found that treatment with BFA (at a concentration of 0.5 μg/ml or higher) reduced by more than 100-fold the viral yield (Fig 1A); this observation is in agreement with a previous report (21).

**FIG. 1.**
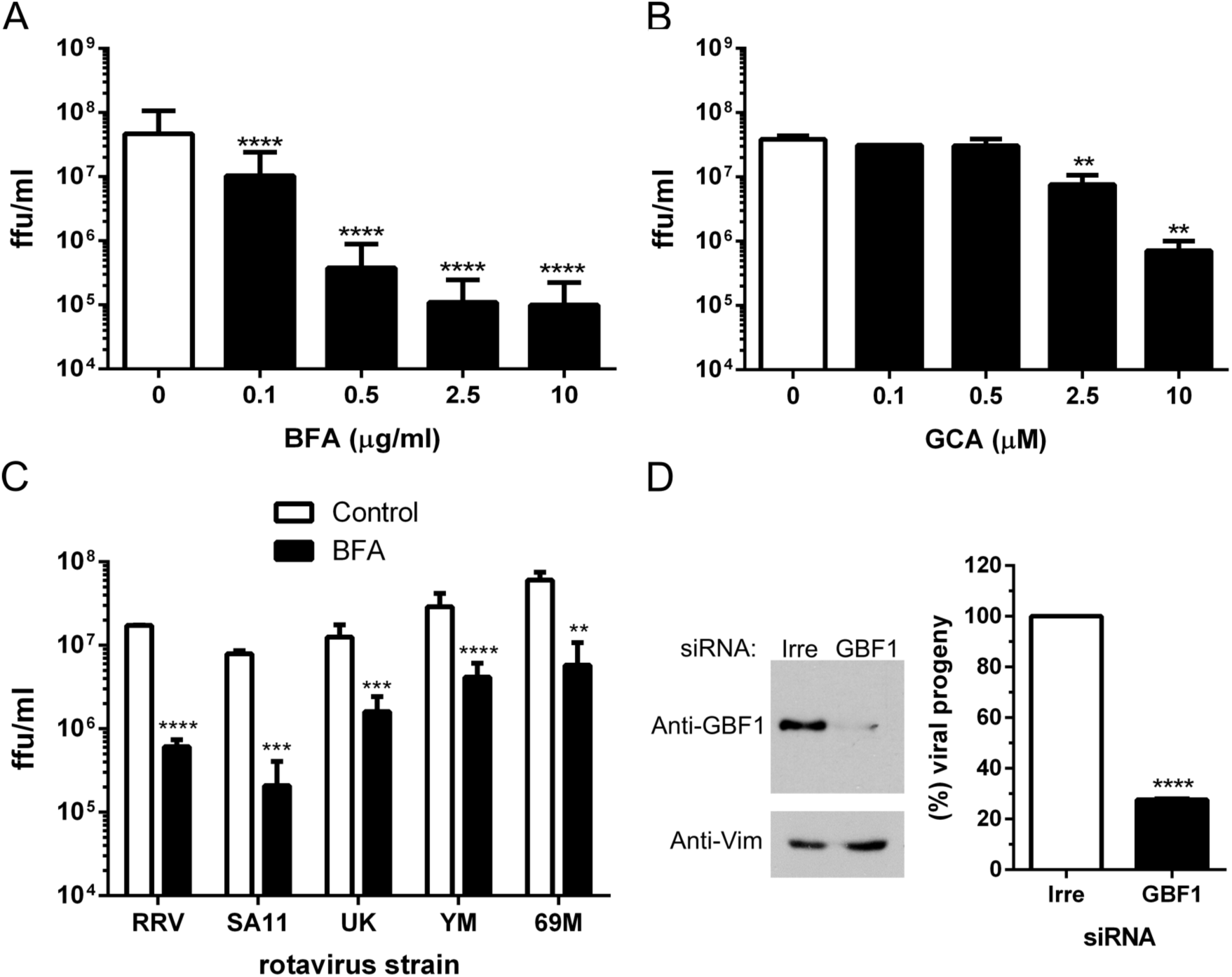
Inhibition of COPI/Arf1 activity reduces the production of viral progeny. MA104 cells were pre-treated for 30 min with BFA (A) or GCA (B) at the indicated concentrations. Treated cells were infected with RRV at an MOI of 5 in the presence of the inhibitors for 1h, the unbound virus was removed, and fresh media containing the drugs was added. At 12 hpi the cells were harvested, and the viral titer was determined by an immunoperoxidase assay. The arithmetic means ± standard deviations from three independent experiments performed in duplicate are shown. (C) MA104 cells were infected with the indicated rotavirus strains (MOI of 5) in the presence of BFA (0.5µg/ml), and the viral titer (ffu/ml) was determined by an immunoperoxidase assay. The arithmetic means ± standard deviations from two independent experiments performed in duplicate are shown. (D) MA104 cells were transfected with a siRNA to GBF1. Left panel, representative western blot analysis of cells transfected with the indicated siRNA. The expression of GBF1 was detected with a specific antibody. Vimentin (Vim) was used as a loading control. Right panel, at 72 hpt cells transfected with either a scrambled or GBF1 siRNAs were infected with RRV (MOI=5). At 12 hpi, cells were collected and the viral titer was determined. Data represent the percentages of virus progeny, where the cells transfected with an irrelevant siRNA (Irre), correspond to 100% infectivity. The arithmetic means ± standard deviations from three independent experiments performed in duplicate are shown. **=P<0.01,***=P<0.001, ****=P<0.0001.

Similarly, treatment with GCA diminished viral progeny about 50-fold (Fig 1B). Cell treatment with the inhibitors did not alter cell viability at any of the concentrations tested, as determined by an LDH assay (data not shown). Moreover, the reduction of viral yield was independent of the rotavirus strain tested, since BFA also significantly inhibited the replication of SA11 (simian origin), UK (bovine origin), 69M (human origin), and YM (porcine origin) (Fig 1C). However, the extent of inhibition differed among the strains; while the replication of SA11 was reduced to a level similar to RRV (about 100-fold), that of rotavirus strains UK, 69M, and YM was reduced by about 10-fold.

To determine whether GBF1 was directly involved in virus replication, we transfected MA104 cells with an siRNA to GBF1, which very efficiently knocked-down the synthesis of this protein (Fig. 1D, left), and 72 h post-transfection (hpt), cells were infected with RRV. After 12 hpi the cells were harvested, and the viral yield was determined. Silencing the expression of GBF1 reduced the yield of viral progeny by about 70% as compared to that produced in cells transfected with control, irrelevant siRNA (Fig 1D, right). These results, together with the fact that GCA decreases RRV replication to a level similar to that observed with BFA, strongly suggest that GBF1 activity might be involved in the replication cycle of rotaviruses.

### BFA and GCA inhibit a post-entry stage of the virus replication cycle

To evaluate the step of the virus lifecycle affected by BFA and GCA, the drugs were added at different times post-infection to cells infected with RRV, as indicated in Fig. 2A, and at 12 hpi the cells were lysed, and the viral yield was determined. The addition of the drugs at 0 and 2 hpi decreased the viral yield by more than 90%; the inhibitory effect was less evident at subsequent times post-infection. When added at 4 hpi, the drugs reduced only around 60% of the viral progeny production, and at later times they showed no significant inhibitory activity (Fig 2A). The fact that the drugs showed a similar inhibitory effect when pre-incubated with cells or when added 2 hpi suggests that the inhibitors are most likely affecting the replication of RRV at a post-entry step. These findings are in agreement with the previous observation that the effect of silencing the expression of β-COP on virus replication was not relieved by transfection of RRV double-layered particles into the siRNA-treated cells (19). Of interest, the fact that the effect of the drugs seems to decline when added at 4 hpi, time at which the assembly of virus particles has already started (Fig. 2B), suggests that the inhibitors could interfere with the virus assembly process. The similar effect observed for GCA and BFA supports the initial observation that the relevant target for the drugs is GBF1.

**FIG. 2.**
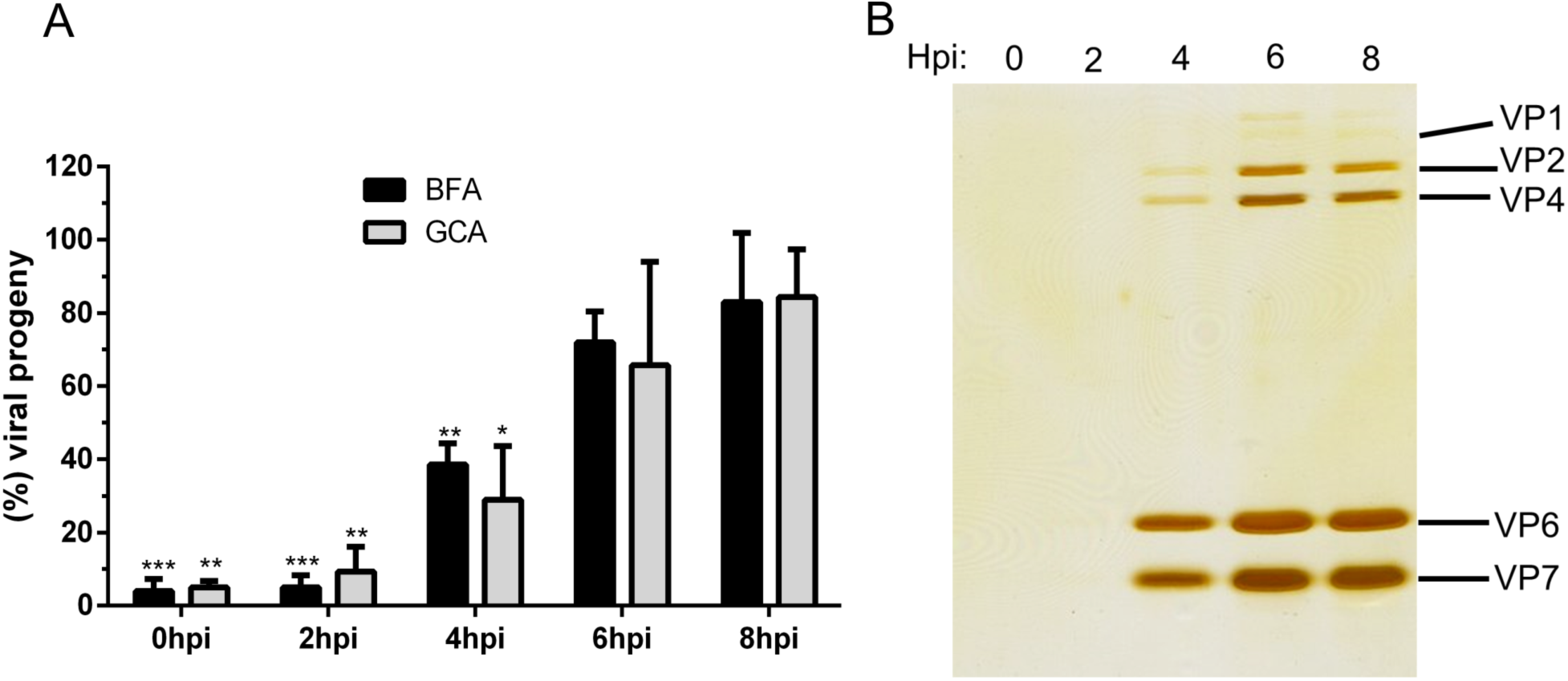
BFA and GCA inhibit rotavirus replication at a post-entry step. (A) MA104 cells were infected with RRV (MOI=5) at 37 °C for 1h. Unbound virus was removed, and 0.5 μg/ml of BFA or 10 μM of GCA were added at the indicated times post-infection. At 12hpi, the cells were lysed, and the viral titer was determined. Cells Data represent the percentage of the virus progeny obtained from untreated cells that correspond to 100% infectivity. The arithmetic means ± standard deviations from two independent experiments performed in duplicate are shown. *=P<0.1, **=P<0.01, and ***=P<0.001. (B) Representative gel electrophoresis analysis of RRV particles semi-purified at different times post-infection. The same proportion of each sample was loaded onto the gel, which was silver stained. The migration pattern of the viral structural proteins is indicated.

### BFA and GCA block the production of TLPs

To confirm that the decrease in viral yield was related to a defect in virus assembly, MA104 cells were infected with RRV in the presence of the drugs and the assembly of viral particles was analyzed by CsCl density gradients. In control conditions, two opalescent bands were observed in the gradients (Figs. 3A and 3C), which were shown to correspond to TLPs and DLPs by PAGE analysis (Figs. 3B and 3D). In contrast, in cells treated with BFA or GCA, a single major band that migrated at a density similar to that of DLPs was detected, and by PAGE was shown to contain particles formed exclusively by proteins VP1, VP2 and VP6 (Fig. 3B). These results indicate that the inhibitors block the assembly of the outermost protein layer, formed by VP4 and VP7, onto the correctly assembled DLPs.

**FIG. 3.**
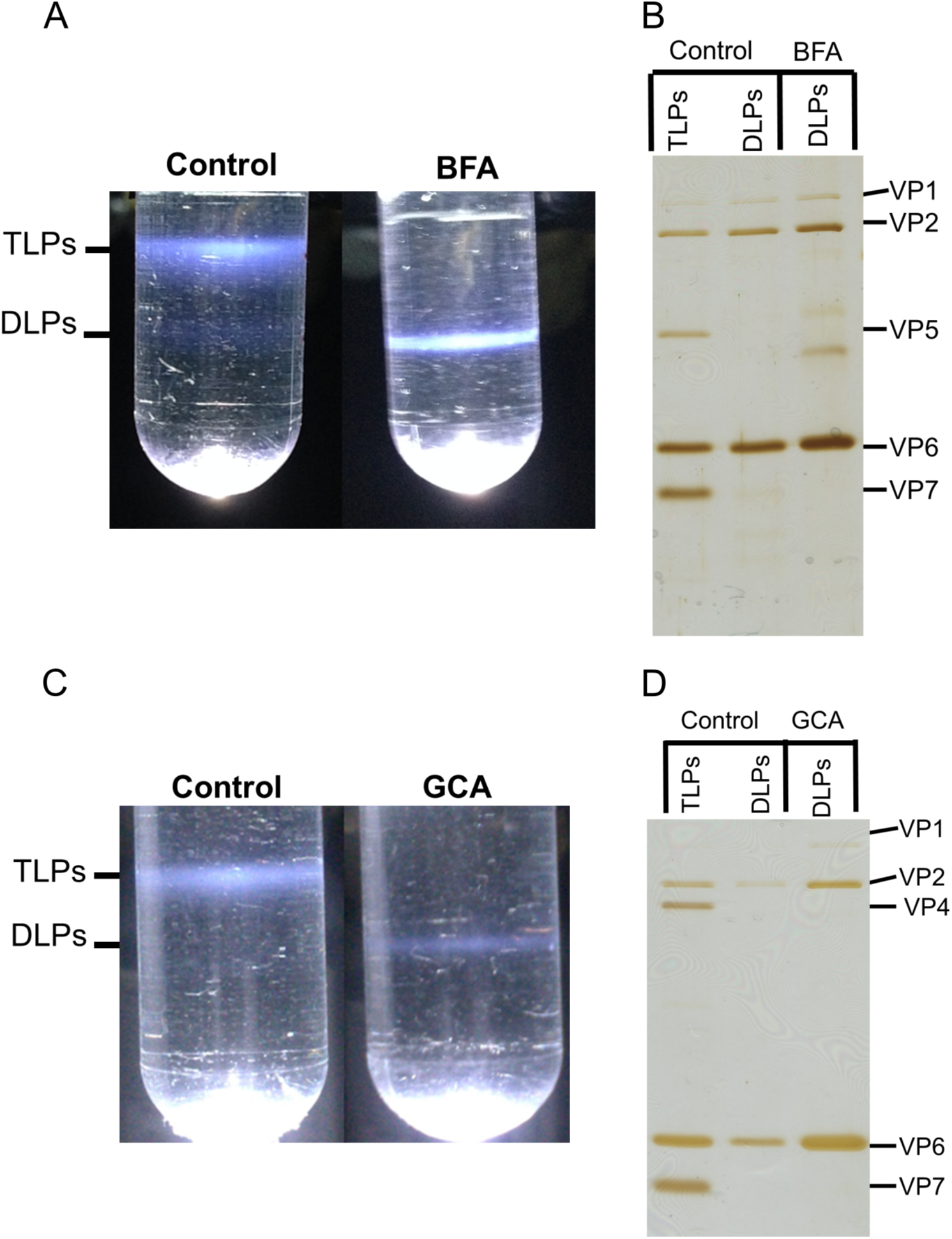
BFA and GCA block the production of TLPs. (A) Isopycnic CsCl gradients of viral particles assembled in the absence or presence of 0.5µg/ml BFA (A) or 10µM GCA (C). (B) and (D) Gel electrophoresis analysis of the viral proteins in the bands detected in the isopycnic gradients shown in panels A and C. The same proportion of each collected band was loaded onto the gel, which was silver stained. The migration of the viral structural proteins in the gels is indicated.

These results were confirmed by negative-staining electron microscopy analysis of drug-treated infected cells. For this, MA104 cells were infected with RRV in the presence of the drugs, and 6 hpi cells were fixed and processed for electron microscopy. In control, untreated cells, typical electro-dense viroplasm structures were observed near the ER membrane. Furthermore, viral particles, presumably DLPs, appear to be budding into the lumen of the ER leading to the formation of membrane-enveloped particles. Many of the viral particles within the ER seem to have lost their lipid envelope and look like mature TLPs (Fig 4A). We found that neither BFA nor GCA prevented the budding of DLPs into the ER to form membrane-enveloped particles; however, the assembly pathway seems to be arrested at this point since the intermediate enveloped particles accumulated within the ER and they did not seem to mature to TLPs (Fig. 4B and 4C). It should be noted that the intermediate membrane-enveloped particles are not detected in the CsCl gradients since they most probably lose their membrane upon with trichloromonofluoromethane extraction, and are rather detected as DLPs.

**FIG. 4.**
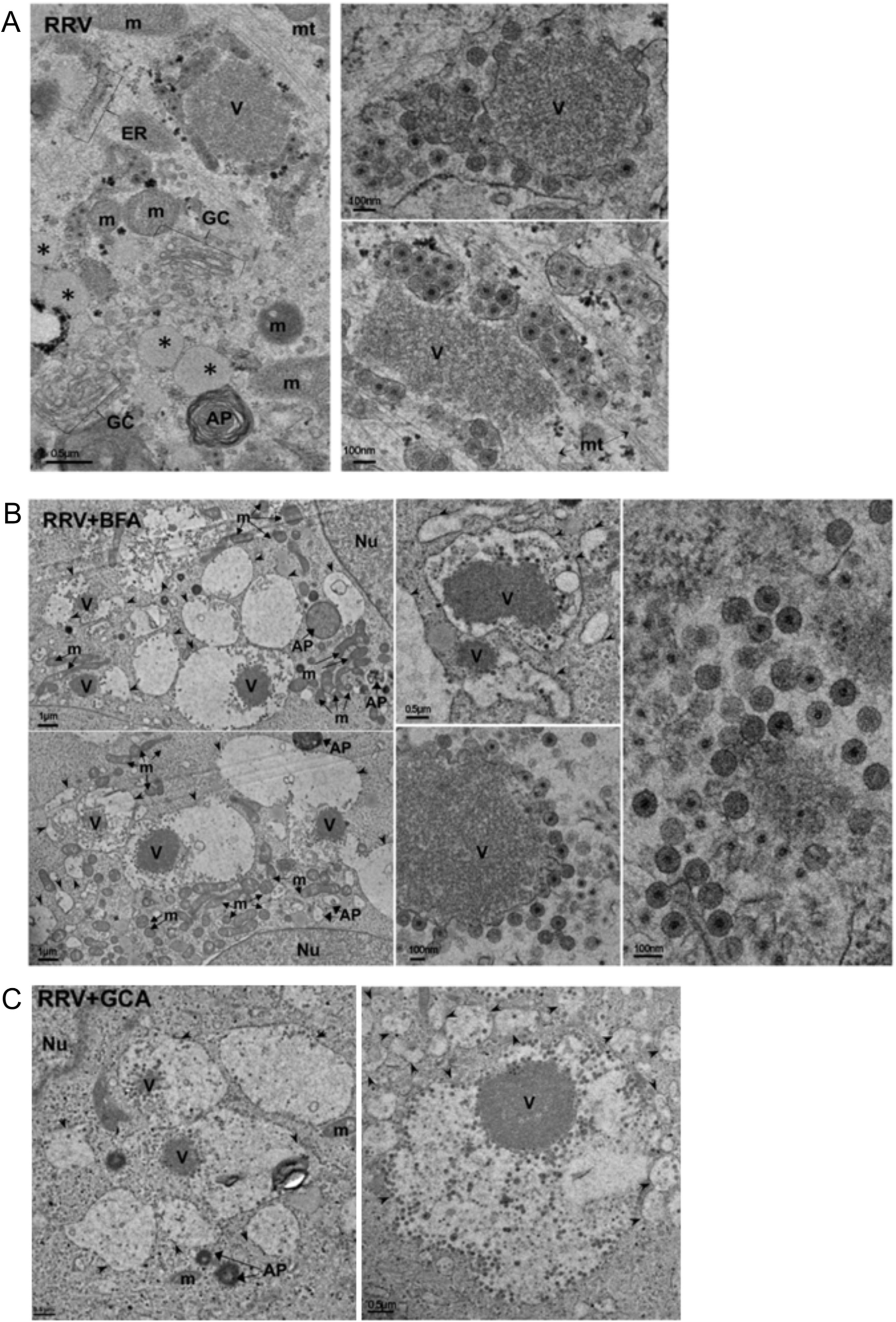
BFA and GCA do not block the budding of rotavirus DLPs into the ER. MA104 cells untreated (A) or treated with 0.5µg/ml BFA (B) or 10µM GCA (C) were infected with RRV at MOI of 250 VFU per cell. At 6hpi, the cells were fixed and processed for thin section electron microscopy. Scale bar is 100 nm. The arrowheads indicate the swollen vesicles of the ER observed in the presence of BFA or GCA. Nu, nucleus; V, viroplasm; m, mitochondria; AP, autophagosome; mt, microtubules; *, vacuoles; ER, endoplasmatic reticulum and GC, Golgi complex.

### The electrophoretic mobility of VP7 and NSP4 is altered during the inhibition of GBF1 activity

Since it has been reported that BFA alters the electrophoretic mobility of VP7 and NSP4 (21), we decided to investigate if the effect of GCA in virus assembly was related to a modification of these viral proteins. MA104 cells were infected with RRV in the presence of the inhibitors as described above, and at 7.5 hpi, the cellular proteins were pulse-labeled with ^35^S-Met for 30 min and chased for 2 h. As seen in Fig. 5A, the inhibitors did not affect the synthesis of either cellular or viral proteins. However, the viral glycoproteins VP7 and NSP4 had modified electrophoretic mobility in the presence of the drugs. In treated cells, VP7 migrated slower than in control cells, while NSP4 showed faster mobility. This distinct electrophoretic behavior was more clearly observed in drug-treated infected cells labeled with ^3^H-mannose, conditions in which only the two viral glycoproteins are labeled (Fig. 5B).

**FIG. 5.**
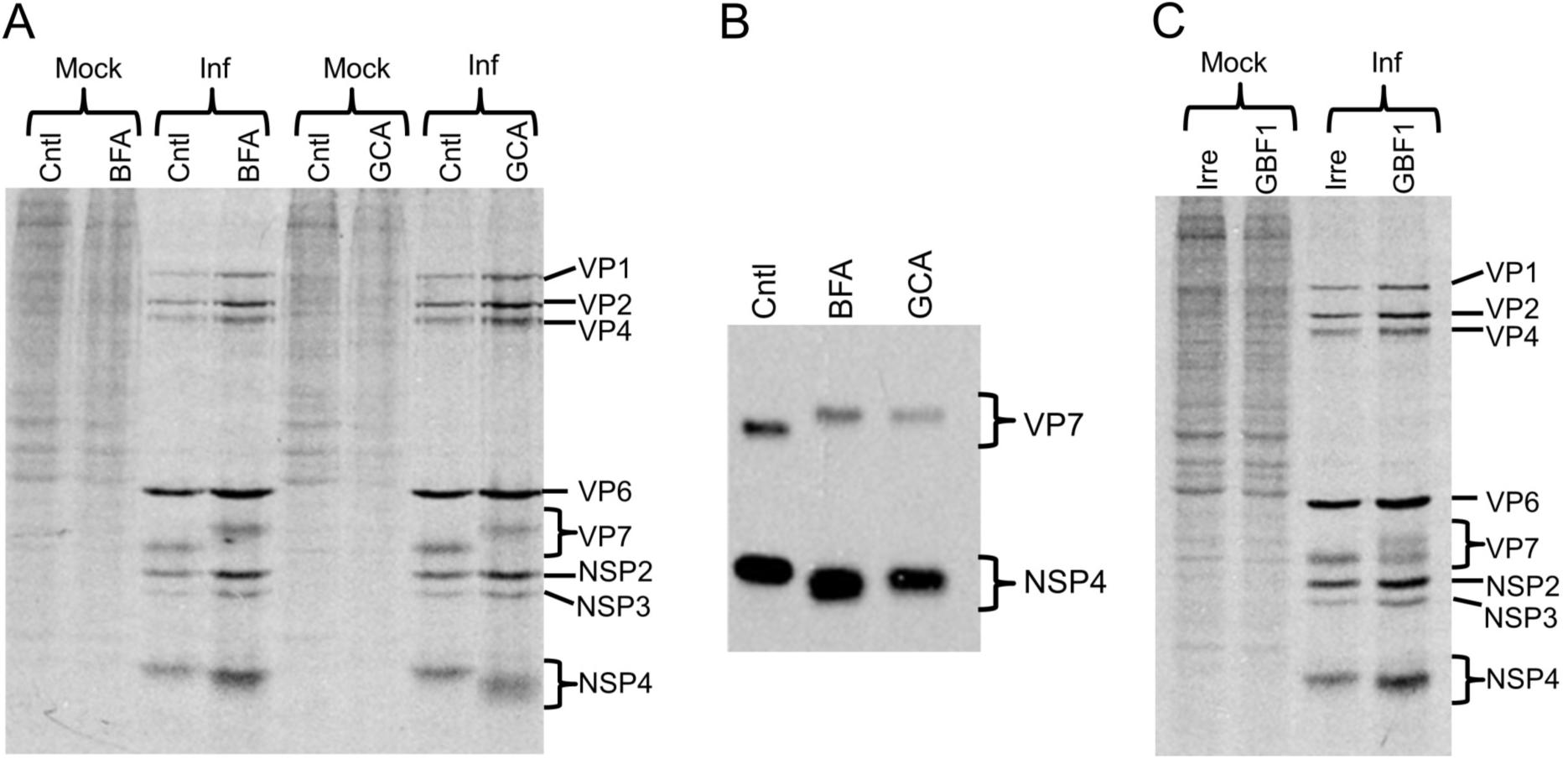
Inhibition of the GBF1 activity changes the electrophoretic mobility of VP7 and NSP4. (A) Autoradiography of mock infected or infected lysates (RRV, MOI of 5) in the presence of BFA (0.5µg/ml) or GCA (10µM). At 7.5 hpi, the cells were pulse-labeled with 25 µCi/ml of ^35^S-Met/Cys for 30 min and harvested at 10 hpi. (B) Fluorography of infected lysates in the presence of BFA (0.5µg/ml) or GCA (10µM). At 6.5 hpi, the cells were radiolabeled with 200 µCi/ml of ^3^H-mannose for 1.5 h and harvested at 10 hpi. (C) Autoradiography of lysates transfected with the indicated siRNA and mock-infected or infected with RRV (MOI=5). At 7.5 hpi, the cells were pulse-labeled with 25 µCi/ml of ^35^S-Met/Cys. The migration pattern of the viral structural and non-structural proteins in the gels is indicated.

To confirm that the alterations in the electrophoretic mobility of VP7 and NSP4 were due to the inhibition of GBF1, MA104 cells were transfected with the siRNA to GBF1 to silence its expression, and infected with RRV; at 7.5 hpi the cells were pulse-labeled with ^35^S-Met as described above. As observed when the pharmacological inhibitors were used, knocking down the expression of GBF1 also caused changes in the electrophoretic mobility of VP7 and NSP4 (Fig 5C); in this case, the change was partial, most likely due to incomplete GBF1 knock down in all cells. To our knowledge GBF1 is the first identified regulatory cellular factor that affects the mobility of VP7 and NSP4.

Furthermore, we also observed that the modification of the electrophoretic mobility of VP7 and NSP4 was independent of the rotavirus strain, since these changes were also observed for rotavirus strains SA11, UK, 69M, and YM, after BFA treatment (data not shown). These results indicate that despite the different amino acid sequences of VP7 and NSP4 among the various rotavirus strains, the lack of GBF1 activity inhibits a common posttranslational processing.

### Modification of the electrophoretic mobility of VP7 in the presence of BFA or GCA is not related to changes in the structure of its carbohydrate chain

It has been previously shown that the treatment of rotavirus-infected MA104 cells with BFA reduces the electrophoretic mobility of VP7, and it was proposed that this change could be related to an altered oligosaccharide chain (21). To determine if the abnormal migration of VP7 in the presence of the inhibitors was due to a modification of its carbohydrate moiety, we characterized the effect of the drugs on the migration of the VP7 protein and replication of the mutant strain SA11-CL28, a simian rotavirus SA11 variant that lacks the N-glycosylation site in VP7 and thus is non-glycosylated (51). The absence of glycosylation of this protein was confirmed by metabolically labeling the virus with ^3^H-mannose. NSP4, but not VP7 was labeled, as detected by PAGE and fluorography (data not shown).

Similar to that observed with wild type SA11 (SA11wt), treatment of cells with BFA or GCA decreased yield of mutant SA11-CL28 virus by more than 90% (Fig. 6A). The drug treatments also reduced the electrophoretic mobility of both, the glycosylated VP7 of SA11wt and the non-glycosylated VP7 of the SA11-CL28 (Fig 6B), suggesting that the change in mobility of VP7 in cells treated with the drugs is not due to a change in glycosylation. Furthermore, by characterizing the electrophoretic mobility of a recombinant RRV VP7 protein overexpressed in MA104 cells, we show that the change observed after BFA treatment is not due to either N- or O-glycosylation or to phosphorylation (Fig. S1). In addition, a recombinant RRV VP7 protein with the N-glycosylation site mutated (69-NST-71 to 69-QSG-71) still showed a reduced electrophoretic mobility after treatment with BFA or GCA, confirming the results observed with the mutant SA11-CL28 virus (Fig. S2).

**FIG. 6.**
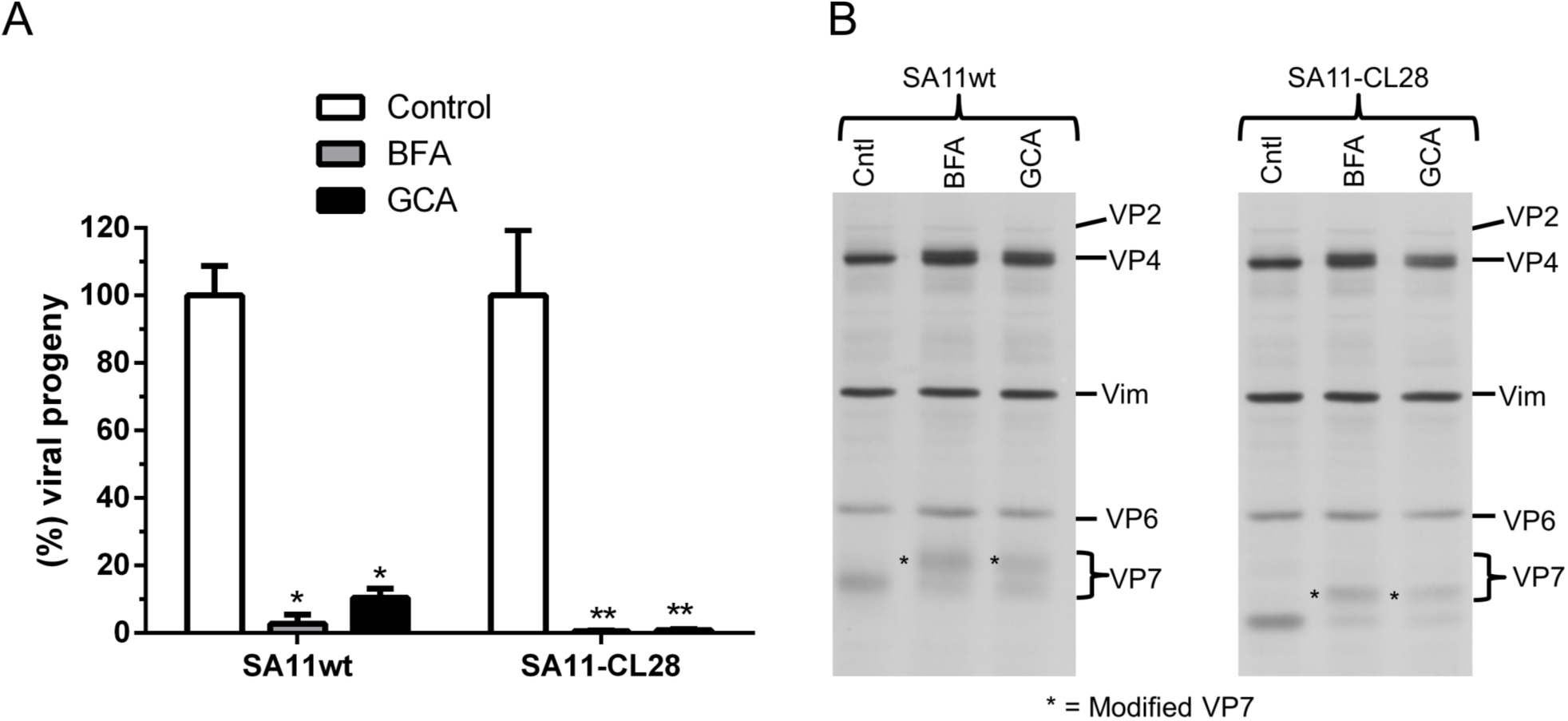
The alteration of the electrophoretic mobility of VP7 in the presence of BFA or GCA is not related to modified glycosylation. (A) MA104 cells were pre-treated for 30 min with BFA (0.5 µg/ml) or GCA (10 µM), and then, infected with wild type SA11 (SA11wt) or the mutant SA11-CL28 strain, both at an MOI of 5, in the presence of inhibitors for 1 h. Unbound virus was removed, and fresh media containing BFA or GCA was added. At 12 hpi, the cells were harvested, and the viral titer was determined. Data represent the virus progeny relative to untreated cells. The arithmetic means ± standard deviations from two independent experiments performed in duplicate are shown. *=P<0.1, **=P<0.01. (B) Representative western blot analysis of proteins from untreated (control), BFA- or GCA-treated cells infected with SA11wt or SA11-CL28 at 8hpi. Expression of structural viral proteins were detected with an anti-rotavirus polyclonal serum raised against purified TLPs. Vimentin (Vim) was used as a loading control. The migration pattern of the viral structural proteins in the gels is indicated. The asterisks mark the position of the slow-migrating VP7 protein (modified VP7).

### An inefficient trimerization of VP7 may be responsible for the defective assembly of TLPs

It has been shown that in the absence of VP7 the transiently enveloped intermediate particles do not mature to infectious virus (52); it was proposed that after budding into the lumen of the ER, VP7 assembles into the DLPs excluding the lipid envelope during this process as well as NSP4, yielding the mature infectious virions (52). Since VP7 exists as trimers in the mature, infectious virus particles (17), we evaluated whether BFA and GCA treatments affected trimerization of the protein. For this, MA104 cells were infected with RRV in the presence of either inhibitor, and 6 hpi the cells were fixed and co-stained with an antibody to the viral nonstructural protein NSP2 that forms part of the viroplasms, and with monoclonal antibody 159 (MAb 159) to VP7, which only recognizes the trimeric form of the protein (18, 53).

In rotavirus-infected, untreated cells, the trimeric isoform of VP7 was observed in the cytoplasm of infected cells, surrounding the viroplasms (Figure 7A). In contrast, in cells treated with BFA or GCA, even though the presence of NSP2 was detected, no signal of the trimeric form of VP7 was observed (Fig 7A). To verify that the lack of trimers was not due to the absence of VP7, we conducted the same assay but using mAb M60, which primarily recognizes the monomeric, non-virion-associated form of VP7 (18, 53). In the absence of inhibitors, the monomeric form of VP7 showed a reticular pattern distributed across the cell cytoplasm, with some of the protein surrounding viroplasms (Figure 7A). Treatment with the inhibitors did not abolish the signal of the VP7 monomers, although the distribution of the protein changed to a punctate pattern (Fig. 7A). Taken together, the treatment with BFA or GCA prevents the formation of VP7 trimers. Of interest, the block in the trimerization of VP7 seems to be independent of alterations in its glycosylation pattern, since BFA also inhibited the trimerization of the VP7 protein of SA11wt and SA11-CL28 viruses without apparently altering the intracellular levels of the monomeric form of the protein (Fig 7B and 7C).

**FIG. 7.**
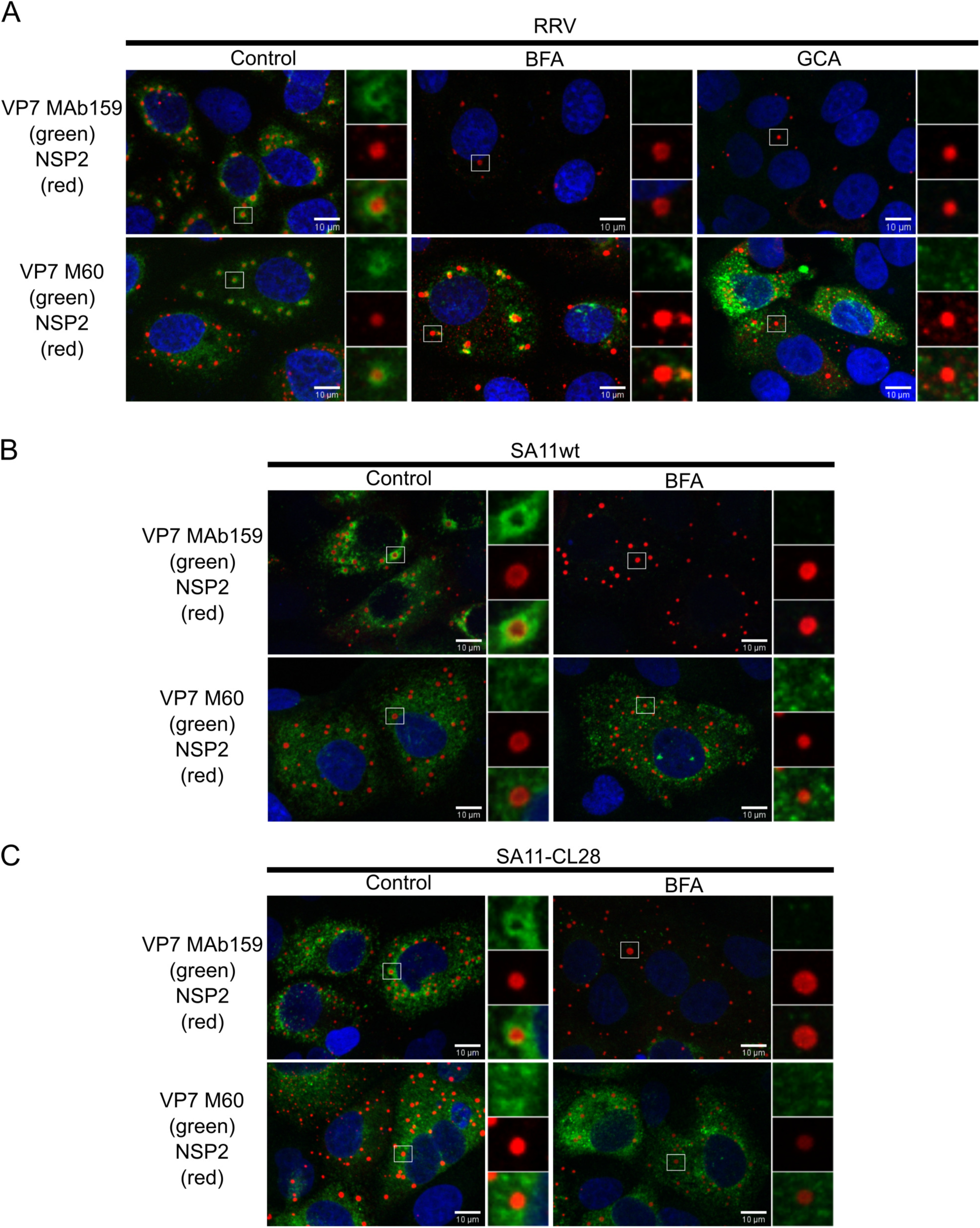
BFA and GCA inhibit the trimerization of the glycosylated and non-glycosylated forms of VP7. Immunofluorescence of MA104 cells infected with RRV (A), SA11wt (B), or SA11-CL28 (C) at an MOI of 1 in the absence or presence of BFA (0.5 μg/ml) or GCA (10 μM). At 6 hpi, the cells were fixed and co-immunostained with either antibodies against NSP2 (red) and VP7 trimers (MAb159, green) (upper row) or NSP2 (red) and VP7 monomers (M60, green) (lower row). The nuclei (blue) were stained with DAPI. The open box corresponds to amplified images at the right column of each image. Scale bar is 10 µm.

We also found that knocking down the expression of GBF1 abolished the trimerization of VP7 (Fig. 8); similar to the experiments with the pharmacological inhibitors, the trimeric form of VP7 (MAb 159) was detected at 6 hpi in the majority of the infected cells transfected with an irrelevant siRNA. In contrast, in cells transfected with the siRNA to GBF1, the trimeric form of VP7 could not be detected, even though the cells were positive for NSP2. However, as described above, the absence of trimers was not due to a deficient synthesis of VP7, since in GBF1-silenced cells the monomeric form of VP7 (detected by MAb M60), was observed in most cells (Fig 8).

**FIG. 8.**
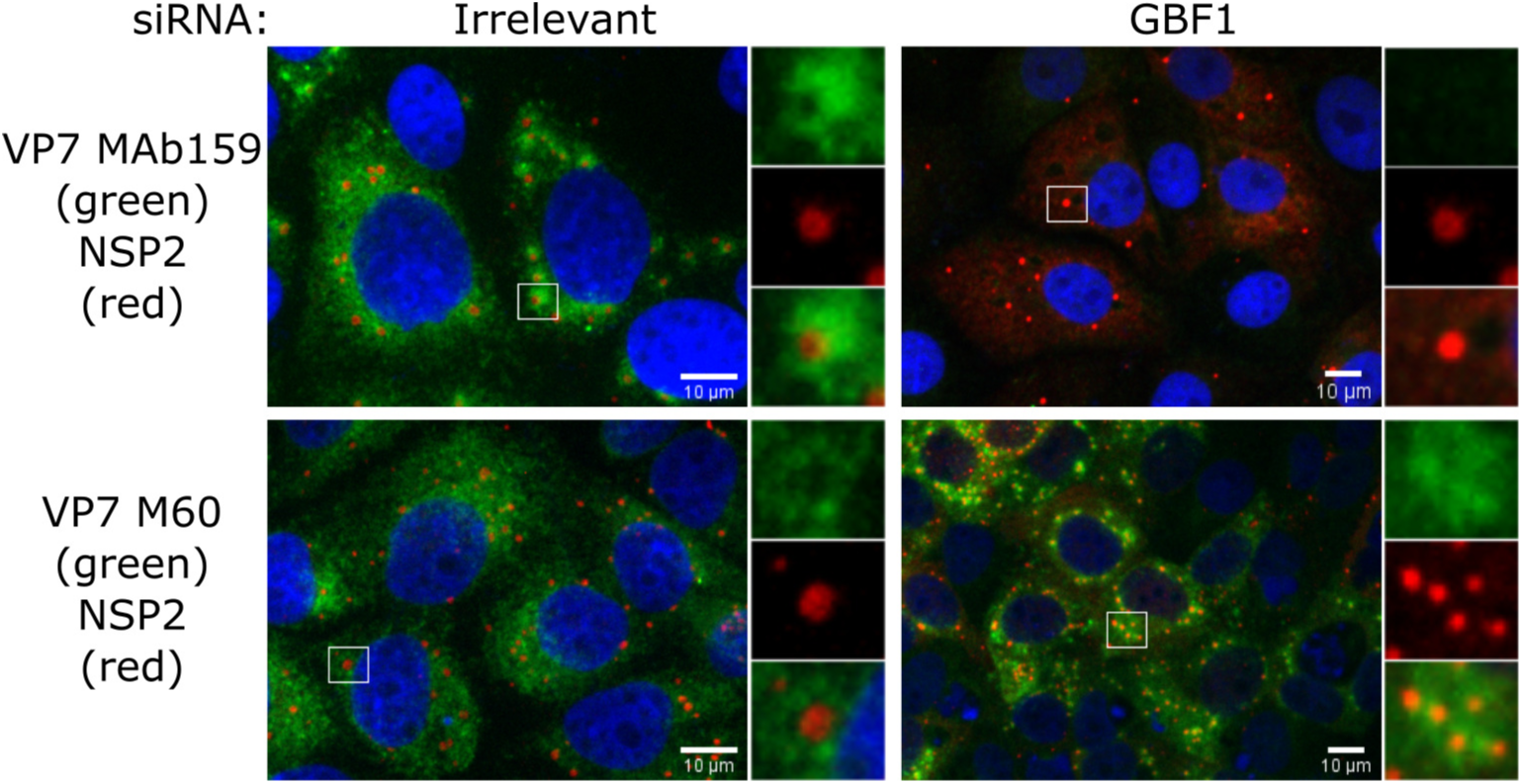
Silencing the expression of GBF1 blocks VP7 trimerization. Immunofluorescence of MA104 cells transfected with the indicated siRNA and at 72 hpt, cells were infected with RRV (MOI, 3). At 6 hpi, the cells were fixed and co-immunostained with antibodies against either NSP2 (red) and VP7 trimers (Mab159, green) (upper panel) or NSP2 (red) and VP7 monomers (M60, green) (lower panel). The cells nuclei (blue) were stained with DAPI. The open box corresponds to magnified images shown at right. Scale bar is 10 µm.

### NSP4 is involved in VP7 trimerization

To explore if the lack of trimerization of VP7 could be related to the change in electrophoretic mobility of NSP4, the expression of NSP4 was silenced by RNAi, and the trimerization of VP7 was assessed in these cells by immunofluorescence microscopy. The efficiency of the knockdown of NSP4 was verified by Western blot (Fig. 9B, upper panel). The presence of VP7 in trimeric or monomeric form was determined in infected cells that were transfected with the siRNA to NSP4 or with an irrelevant siRNA and were stained with MAbs 60 or 159. As above, cells transfected with an irrelevant siRNA contained the trimeric form of VP7. In contrast, in cells transfected with the siRNA to NSP4, a larger proportion of infected cells did not contain the trimeric form of VP7, even though they were infected as judged by the presence of NSP2 (Fig. 9A). Quantification of these cells showed that in control conditions ∼75% of the infected cells were positive for MAb 159 staining, while in cells in which the expression of NSP4 was knocked down, only 40% of the infected cells showed some signal for the trimeric form of the protein (Fig. 9B, lower panel). Although not statistically significant, this difference suggests a tendency of VP7 to form trimers when NSP4 is present, suggesting that NSP4 facilitates or participates in the trimerization of VP7. These results are consistent with a previous report showing that NSP4 might stimulate the assembly of VP7 (54).

**FIG. 9.**
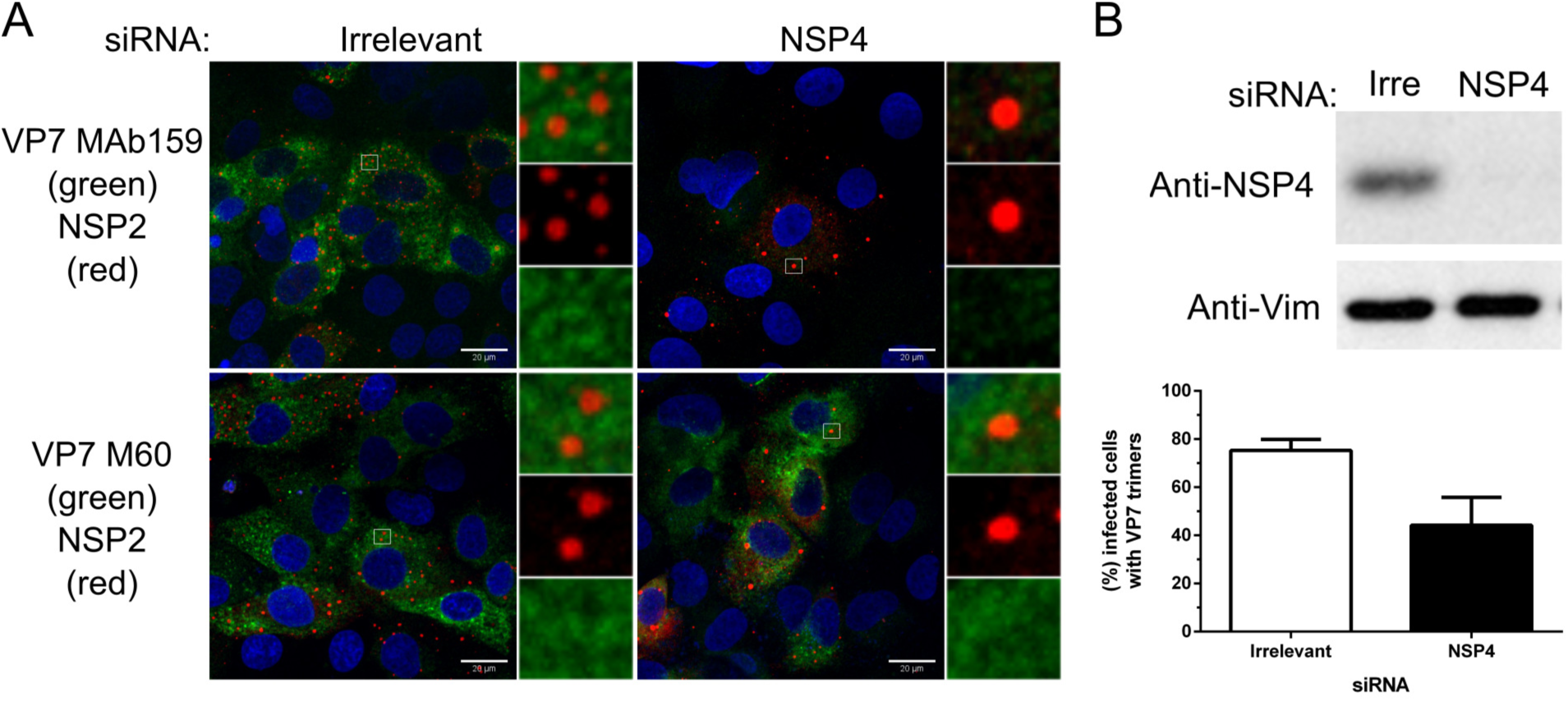
Silencing the expression of NSP4 inhibits the trimerization of VP7. (A) Immunofluorescence of MA104 cells transfected with the indicated siRNA, and at 72 h post-transfection, the cells were infected with RRV (MOI, 3). At 6 hpi, cells were fixed and co-immunostained with antibodies against either NSP2 (red) and VP7 trimers (MAb159, green) (upper row) or NSP2 (red) and VP7 monomers (M60, green) (lower panel). Cells nuclei (blue) were stained with DAPI. The open box indicates magnified images at the right of each image. Scale bar is 20 µm. (B) Upper panel, representative western blot analysis of cells transfected with the indicated siRNA. The expression of NSP4 was detected with a specific antibody. Vimentin (Vim) was used as a loading control. Lower panel, MA104 cells transfected with the indicated siRNA were infected with RRV and processed for immunofluorescence as described in (A). The number of infected cells positive to VP7 trimers (green cells) was determined by counting 1000 cells in each condition. Values are expressed as a percentage of the total infected cells. The arithmetic means ± standard deviations from three independent experiments are shown.

To further explore whether the trimerization of VP7 depends on the presence of NSP4, Hek293 cells expressing constitutively the T7 polymerase (T7pol) (55) were co-transfected with plasmids D1R and D12L encoding the two subunits of the vaccinia virus capping enzyme, which efficiently caps the T7pol RNA polymerase transcripts in the cytoplasm (56), and with plasmids encoding the non-glycosylated forms of VP7 or NSP4, or with both plasmids simultaneously. At 24 hpt the cells were fixed and immunostained for NSP4 and VP7. In cells in which the plasmid for VP7 was transfected alone, the monomeric form of VP7 (MAb M60) was clearly visible, whereas there was no signal with MAb 159 that detect the trimer; in contrast, when the plasmids for VP7 and NSP4 were co-transfected, trimeric VP7 was detected and colocalized with NSP4 (Fig. 10). These results confirm that the presence of NSP4 is relevant (directly or indirectly) for the correct assembly of VP7 into trimers.

**FIG. 10.**
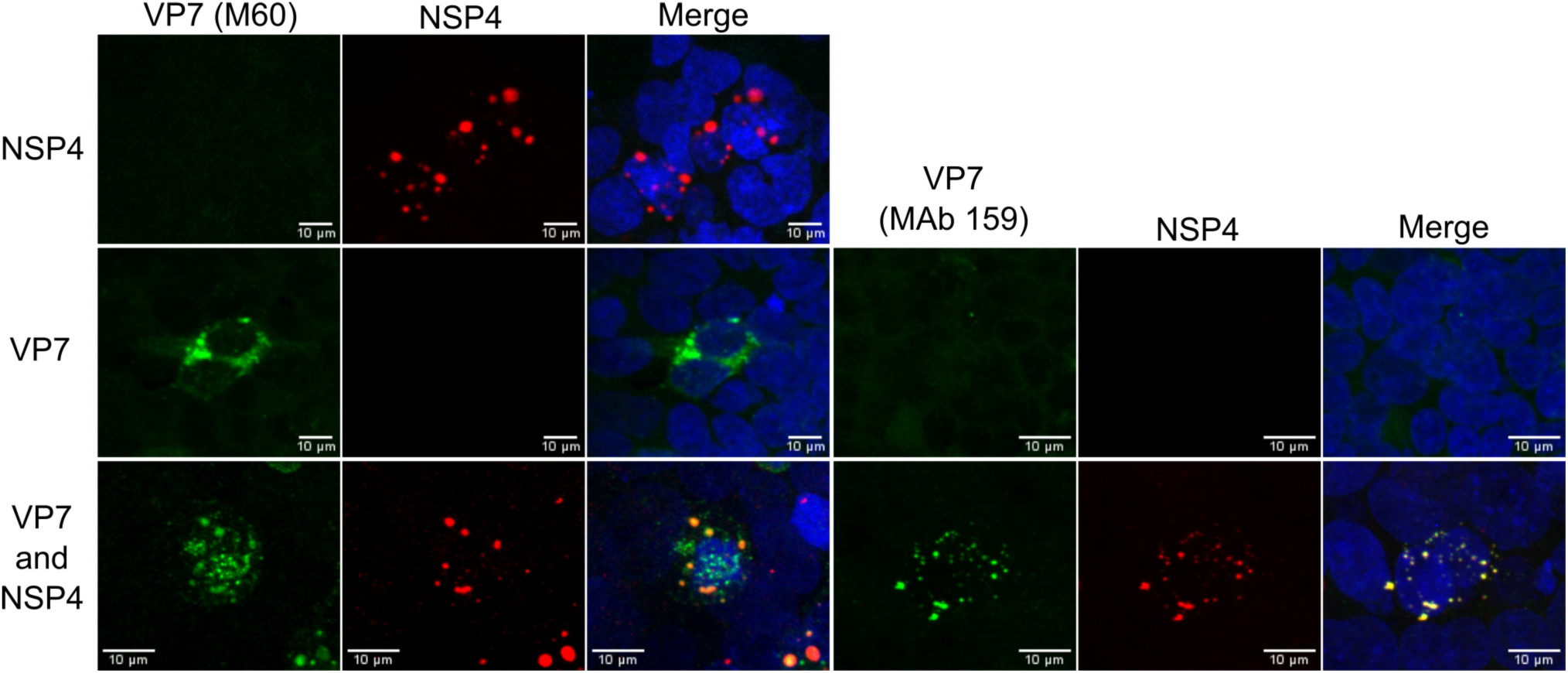
NSP4 facilitates the VP7 trimerization. Immunofluorescence of Hek293-T7 RNA polymerase cells co-transfected with the VV capping enzyme expression plasmids (D1R and D12L) and either a plasmid encoding for non-glycosylated VP7, NSP4, or a combination of both plasmids. At 24 hpt, the cells were fixed and co-immunostained with antibodies directed either to NSP4 (red) and VP7 monomers (M60, green) or NSP4 (red) and VP7 trimers (MAb159, green). The cells nuclei (blue) were stained with DAPI. Scale bar is 10 µm.

### Distinct domains of GBF1 are essential for rotavirus replication

To explore the potential involvement of the different GBF1 domains in rotavirus progeny production, we compared the replication of the virus in Hek293 cells transfected with GBF1 construct containing the A795E mutation (GBF1/795) that confers resistance to BFA (48, 57), or with a series of mutant GBF1/795 proteins (Fig 11A) (58).

For these assays, Hek293 cells were transfected with the different constructs, and at 24 hpt, the cells were infected with RRV. At 12 hpi the cells were harvested, and the viral yield was determined. Figure 11B shows that the full-length GBF1/795 was able to rescue the replication of rotavirus in the presence of BFA. In addition, the GBF1/795/1531t was able to rescue, albeit partially, viral replication. In contrast, none of the other GBF1 mutants could support virus replication. These findings indicate that the C-terminal domain of GBF1 is not absolutely critical for replication of the virus, while all other domains of GBF1 are required.

We also tested a GBF1/795 mutant that encoded a protein with an amino acid substitution E794K in the Sec7 domain (GBF1/795/E794K) that is inactive in Arf activation (59), and with a construct in which the C-terminal loop of Sec7 domain (EIVMPEE at positions 883–889) was disrupted by the substitution of 7 alanines, producing a GBF1 protein defective in Arf biding and activation (GBF1/795/7A)(60)(Fig 11A). Both, the catalytic inactive GBF1/795/E794K and GBF1/795/7A mutants were unable to support rotavirus replication in the presence of BFA, suggesting that the ability of GBF1 to activate Arf is important for virus progeny production. All GBF1/795 constructs had been shown to be expressed to a similar level, as determined by western blot (57, 58)

The previous results suggested that Arf1 activation might be important for rotavirus replication. This observation was tested by transfecting MA104 cells with an siRNA to Arf1, which very efficiently knocked-down the synthesis of the protein (Fig. 11C, left), and at 72 hpt, infecting the cells with RRV. At 12 hpi the cells were harvested, and the viral yield was determined. Silencing the expression of Arf1 did not affect the yield of viral progeny as compared to that produced in cells transfected with a control, irrelevant siRNA (Fig 11C, right). These results suggest that GBF1 catalytic activity is essential for rotavirus replication, but the activation of Arf1 is not required.

**Fig 11.**
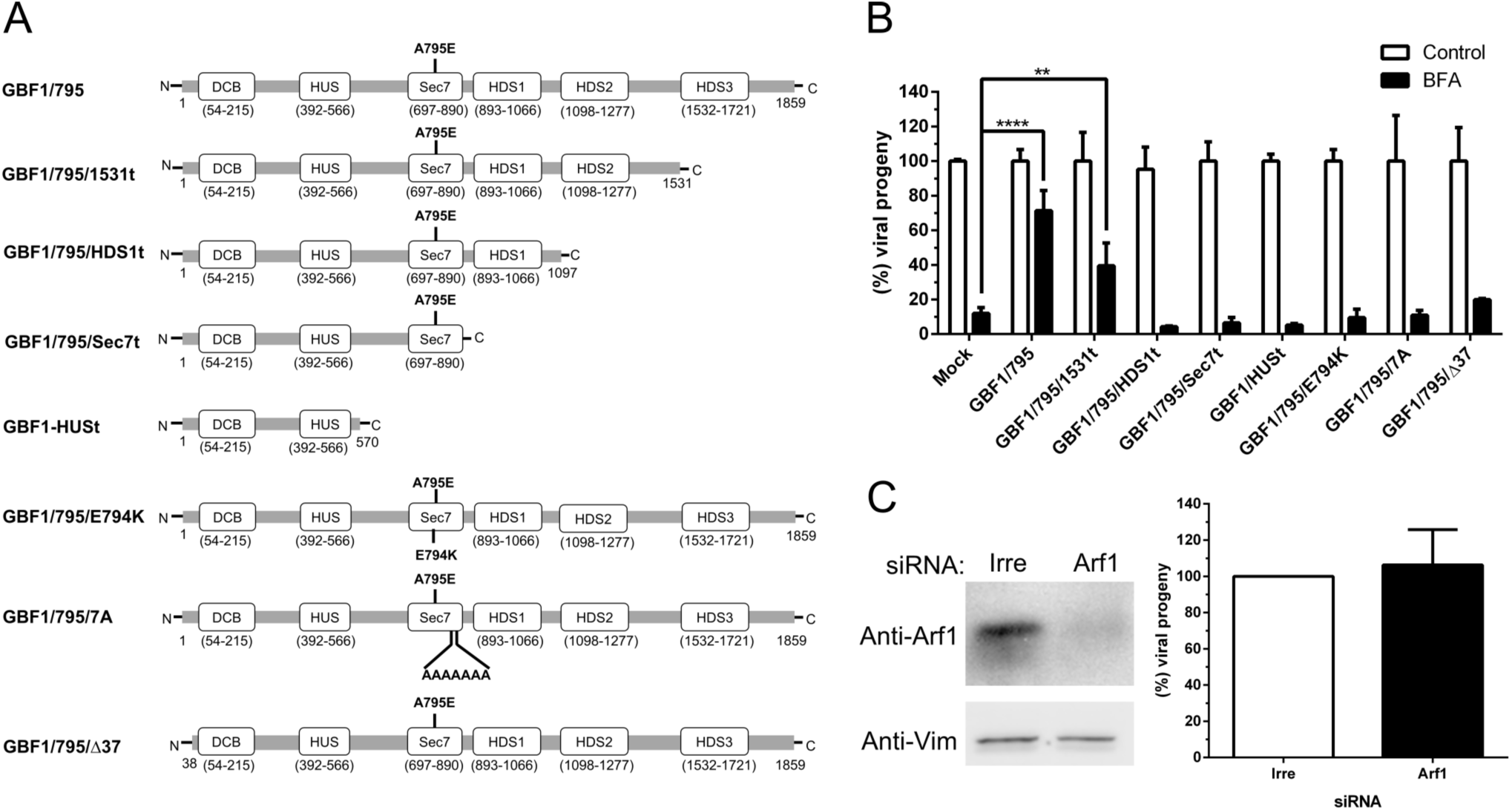
GBF1 catalytic activity but not Arf1 activation is essential for rotavirus replication. (A) Schematic map of GBF1 domain organization and of truncated mutants used in this study, in which the amino acid substitutions are indicated. Numbers in parenthesis indicate GBF1 amino acids. (B) Hek293 cells were transfected with the different GBF1 mutant plasmids, and at 24 hpt cells were pre-treated for 30 min with BFA (0.5 µg/ml), and then, infected with RRV at an MOI of 5, in the presence of BFA for 1 h. Unbound virus was removed, and fresh media containing BFA was added. At 12 hpi, the cells were harvested, and the viral titer was determined. Data represent the virus progeny relative to untreated cells. The arithmetic means ± standard deviations from three independent experiments performed in duplicate are shown. **=P<0.01, ****=P<0.0001. (C) MA104 cells were transfected with a siRNA to Arf1. Left panel, representative western blot analysis of cells transfected with the indicated siRNA. The expression of Arf1 was detected with a specific antibody. Vimentin (Vim) was used as a loading control. Right panel, at 72 hpt scramble siRNA and siArf1cells were infected with RRV (MOI=5). At 12 hpi, cells were collected and the viral titer was determined. Data represent the percentages of virus progeny, where the cells transfected with an irrelevant siRNA (Irre), correspond to 100% infectivity. The arithmetic means ± standard deviations from three independent experiments performed in duplicate are shown.

## DISCUSSION

Eukaryotic cells use elaborated systems to control the traffic of proteins between different organelles. The COPI/Arf1 machinery has been shown to participate in several of these transport processes, including its recent involvement in maturation and function of lipid droplets (LDs) (30–32). The COPI/Arf1 complex is also emerging as an important cellular factor for the replication of several RNA viruses. In the case of influenza (61, 62) and vesicular stomatitis viruses (VSV) (63, 64) the COPI/Arf1 complex seems to play a role in their entry process. Moreover, for VSV as well as for hepatitis C virus (65–67), mouse hepatitis coronavirus (68), chikungunya virus (69), poliovirus (57, 58, 70), coxsackievirus (71) and classical swine fever virus (72), COPI/Arf1 has been shown to be necessary for genome replication as well as for viral protein expression. COPI/Arf1 has also been shown to be critical for viral particle assembly of influenza and the Chandipura virus (62, 73). The specific mechanism through which COPI/Arf1 participate in these processes has not been characterized; however, GBF1, involved in the first step of the COPI/Arf1 transport process, has been shown to play an important role for the replication of some of these viruses. For example, for enteroviruses, the recruitment of GBF1 to the membranous structures where the viral RNA synthesis takes place, mediated by the viral 3A protein, is crucial for genome replication. Moreover, in this case, the function of GBF1 appears independent of Arf1 activation (57, 74).

In this work, we studied the effect of inhibiting the function of the GBF1/COPI/Arf1 machinery on the replication of rotavirus using the pharmacological inhibitors BFA and GCA. We found that both inhibitors induced a significant reduction in the progeny production of different rotavirus strains belonging to different G and P serotypes, indicating that these drugs block a common pathway for all these viruses.

GCA inhibits the COPI/Arf1 machinery activity by selectively inhibiting GBF1; BFA, on the other hand, also inhibits BIG1 and BIG2 (49, 75, 76). BIG1 and BIG2 are involved in vesicular traffic of clathrin-coated vesicles at the TGN (77–79). Our observations suggest that GBF1, and not the BIGs, is the factor targeted by BFA and GCA that causes the reduction of rotavirus replication. The importance of GBF1 was confirmed since knocking down the expression of GBF1 by RNAi decreased the infectivity of RRV by about 70%.

In line with previous observations, we found that in cells treated with BFA and infected with RRV, the electrophoretic mobility of both VP7 and NSP4 was modified, a modification that was proposed to be related to their oligosaccharide chains (21). Since no infectious virus was produced under those conditions, it was also proposed that the altered oligosaccharide processing of VP7 was responsible for the defective assembly of viral particles. Our results indicate that, at least for VP7, the altered electrophoretic mobility is independent of its glycosylation pattern, since both the non-glycosylated VP7 of the SA11-CL28 virus and a recombinant VP7 protein with the N-glycosylation site mutated showed an altered mobility in the presence of the drugs. These results are in agreement with previous findings showing that cell treatment with both BFA and tunicamycin, an inhibitor of N-glycosylation (80), produce a VP7 protein with altered mobility (21). In addition, the changes in VP7 mobility were not due to O-glycosylation or phosphorylation of the protein. The possibility that the drugs block the removal of the signal peptide of VP7 (81) remains to be explored.

The observation that silencing the expression of GBF1 induced changes in the mobility of both VP7 and NSP4 suggests that the activity of this cellular factor is essential for the correct processing of both proteins. Taking into consideration that GBF1 functions at the ER-Golgi interface and that rotaviruses mature in the ER, it seems likely that Golgi-ER transport is the relevant process required for the correct maturation of rotavirus infectious particles. However, considering that LDs have been proposed to play a significant role in the replication cycle of rotaviruses, it remains possible that the protein transport between the ER membrane and LDs is needed for the correct processing of both VP7 (glycosylation independent) and NSP4. However, an unknown function of GBF1 cannot be discarded.

Since the assembly of VP7 onto DLPs has been suggested to require VP7 trimerization, the effect of the drugs on trimerization was evaluated. We found that VP7 was not able to trimerize (as judged by the lack of recognition of VP7 by MAb 159, which specifically interacts with the trimeric but not with the monomeric form of the protein (18)) in the presence of the inhibitors or in cells where the GBF1 expression was silenced,. Our findings differ from previous observations that BFA has minimal effects on the antigenicity of VP7 (as judged by recognition by MAb 159 in immunoperoxidase staining (21)). The reason for this discrepancy might be the methods employed for the detection of the MAb159 signal, immunofluorescence in our work versus immunoperoxidase staining in the previous report (21), which may have different sensitivity.

The drug-induced modification of VP7 could, in principle, prevent its trimerization. However, it is also possible that the modification of NSP4 could impair VP7 trimerization. Even if the NSP4 function as a receptor in the ER membrane is not affected by the drug treatments, it is plausible that the modifications in its glycosylation pattern could affect its role in facilitating (directly or indirectly) the trimerization of VP7 during the removal of the envelope in the last step virus assembly. The role of NSP4 in the trimerization of VP7 is supported by earlier observation where tunicamycin, an inhibitor of N-glycosylation, restricts the maturation of enveloped particles of SA11-CL28 by inhibiting the glycosylation of NSP4 (82). On the other hand, it is known that the stability of the VP7 trimers depends on the presence of calcium (Ca2+) (15, 18), suggesting that alteration in the calcium homeostasis of the cell could be responsible for the lack of the outer layer assembly of rotaviruses. It has been shown that rotavirus infection increases the Ca2+ permeability of the plasma membrane, leading to an enhancement of its intracellular concentration, but that BFA inhibits this enhancement (83). Thus, it cannot be excluded that the deficient trimerization of VP7 is due to altered calcium homeostasis.

The multidomain structure of GBF1 allows it to engage in numerous interactions with other cellular, as well as viral proteins (33, 58, 74). To gain information about the importance of the different domains of GBF1 in rotavirus replication, we tested the ability of different truncated forms of GBF1 to support the production of virus progeny in the presence of BFA. A recombinant full length GBF1 carrying a mutation in the Sec7 domain that renders the protein resistant to the inhibitory action of BFA (GBF1/795) was able to rescue virus replication, as did a mutant GFB1 protein lacking the C-terminal HDS3 domain (albeit less well), suggesting that HDS3 is dispensable for viral replication. Although the precise function of the HDS3 is unknown, it appears essential for GBF1 membrane association, since the GBF1/795/1531t mutant is inefficiently targeted to Golgi, remaining predominantly cytosolic, albeit some fraction of this mutant is able to associate the membranes (unpublished results). It possible that the small fraction of the GBF1/795/1531t mutant that is able to associate to membranes is capable to activate sufficient Arf molecules to allow the lifecycle of the virus to proceed.

The catalytically inactive mutants of GBF1 (GBF1/795/E794K and GBF1/795/7A) failed to support replication of the virus, suggesting that GBF1 activity is essential for rotavirus progeny production. Although the catalytic activity of GBF1 appears essential for rotavirus replication, we found that the activation of Arf1 is not required, since knocking-down the expression of this factor does not affect the production of infectious virus. Thus, it is likely that rotavirus replication requires the activation of other GBF1 substrates, such as Arf4 or Arf5 (Claude et al., 1999; Niu et al., 2005; Szul et al., 2005) that could compensate for the reduced level of Arf1.

Previous findings reported that the most N-terminal 37 amino acids of GBF1 are important for the replication of coxsackie B3 virus and poliovirus (57, 58, 74, 84) by targeting the 3A protein to membranes where the virus replication complexes will assemble. Similarly, we show that this deletion renders GBF1 unable to support rotavirus replication in the presence of BFA. It is possible that this 37 amino acid stretch could bind a rotaviral protein, although it may also interact with a cellular factor to support virus replication. In fact, it has been reported that Rab1b is able to interact with the N-terminus of GBF1 to modulate GBF1 function in the secretory pathway (85). Further experiments will have to be carried out to explore more in detail the mechanism through which GBF1 supports rotavirus replication.

## MATERIALS AND METHODS

### Cell and viruses

MA104 rhesus monkey kidney epithelial cells (ATCC), and Hek293 human embryonic cells stably expressing phage T7 polymerase, kindly provided by Dr. Carlos Sandoval Jaime (Instituto de Biotecnología, Cuernavaca, Mexico) (55), were grown in Dulbecco’s Modified Eagle Medium-Reduced Serum (DMEM-RS) (Thermo Scientific HyClone, Logan, UT) supplemented with 5% heat-inactivated fetal bovine serum (FBS) (Biowest, Kansas City, MO) at 37°C in a 5% CO2 atmosphere. Simian (RRV, G3P5B[3]; SA11, G3P5B[2]; and SA11-CL28, G3P5B[2]), bovine (UK, G6P7[5]), porcine (YM, G11P9[7]), and human (69M, G8P4[10]) rotaviruses were propagated in MA104. Briefly, the viruses were activated with trypsin (10 µg/ml; Gibco, Life Technologies, Carlsbad, CA, USA) for 30 min at 37°C. Afterwards, the activated viruses were adsorbed to the cells for 1 h at 37°C. The unbound virus was removed, and the cells were incubated for 16 h at 37 °C. Finally, the cells were lysed by two cycles of freeze-thawing and the cell debris were removed by centrifugation. The resulting viral lysate was stored at −70°C.

### Reagents and antibodies

Brefeldin A (B7651, Sigma-Aldrich, St. Louis, MO, USA) was dissolved in ethanol, and Golgicide A (345862, Calbiochem, San Diego, CA, USA) was dissolved in dimethyl sulfoxide (DMSO). Monoclonal antibodies (MAbs) to the monomeric (M60) and trimeric (MAb159) forms of VP7 (53) were kindly provided by H. B. Greenberg (Stanford University, Stanford, USA). The rabbit anti-rotavirus polyclonal serum raised against purified TLPs(52), the rabbit polyclonal sera to NSP2(86) and NSP4(87), as well as the rabbit polyclonal sera to human vimentin(52), were produced in our laboratory. Mouse monoclonal antibody to GBF1 was purchased from BD Transduction Laboratories (San Jose, CA, USA). Goat anti-mouse Alexa-488 and goat anti-rabbit Alexa-568-conjugated used as secondary antibodies were obtained from Molecular Probes (Eugene, OR, USA) Horseradish peroxidase-conjugated goat anti-rabbit and anti-mouse antibodies were purchased from PerkinElmer Life Sciences (Boston, MA, USA).

### SiRNAs and plasmids

The small interfering RNA (siRNA) to GBF1, Arf1 and the control siGENOME nontargeting siRNA were purchased from GE Healthcare Dharmacon (Lafayette, CO, USA). For the construction of pcDNA3-VP7 and pcDNA3-NSP4 plasmids, the VP7 and NSP4 genes were cloned from extracts of RRV- and SA11-infected cells, respectively. The cDNA was obtained by RT-PCR using specific primers for amplifying the open reading frame (ORF) of VP7 or NSP4 and cloned into pcDNA3 (Life Technologies, USA) as KpnI-EcoRI fragments. The pcDNA3-non glycosylated VP7 plasmid was obtained by site-direct mutagenesis of the glycosylated VP7 of RRV. Vaccinia virus capping enzyme expression plasmids, pCAG-D1R and pCAG-D12L (Addgene plasmids # 89160 and # 89161, respectively) were a gift from T. Kobayashi (Osaka University, Osaka, Japan). All GBF1 truncations were introduced into the backbone of Venus tagged-GBF1/A795E (described in 57) using QuickChange XL site-directed mutagenesis kit from Agilent Technology. The GBF1/795/E794K construct was generated by introducing the A795E mutation into GFP-tagged GBF1 E794K-GFP construct (described in 59). Following PCR, constructs were transformed into XL 10 gold cells. DNA was isolated and all mutations were confirmed by sequencing. The GBF1/795/7A construct has been described (60). The GBF11/795/Δ37 construct has been described (57).

### Immunoperoxidase assay

MA104 cells were grown in 96-well plates to confluence. The cells were then infected with two-fold serial dilutions of the different viral lysates for 1h at 37°C. After this time, the non-adsorbed virus was removed, and the cells were incubated at 37°C for 16 h. Afterwards, the cells were fixed with 80% acetone in phosphate-buffered saline (PBS) for 30 min at room temperature and then washed twice with PBS. The fixed monolayers were then incubated with a rabbit anti-rotavirus polyclonal serum, followed by incubation with a secondary anti-rabbit antibody conjugated with horseradish peroxidase. Finally, the cells were washed twice with PBS and stained with a solution of 1 mg/ml of carbazole (AEC) in sodium acetate buffer (50 mM, pH 5) with 0.04% H_2_O_2_. The reaction was stopped by washing in tap water, and the infectious focus forming units were counted in a Nikon TMS Inverted Phase Contrast Microscope with a 20X objective.

### Transfection of siRNAs and plasmids

Transfection of siRNAs into MA104 cells was performed with Oligofectamine reagent (Invitrogen, Carlsbad, CA, USA) in 48-well plates using a reverse transfection method as described previously (88). Plasmids were transfected either into MA014 cells or Hek293 cells constitutively expressing the T7 polymerase, using Lipofectamine 3000 (Invitrogen, Carlsbad, CA, USA) according to the manufacturer’s instructions.

### Enrichment of viral particles

RRV (at an MOI of 5) was adsorbed to MA104 cells for 1 h at 4°C to allow the virus to bind to the cells. The unbound virus was removed, and the cells were incubated at 37 °C to allow the infection to proceed. At different times post-infection cells were washed with EGTA (3 mM) for 5s to release any viral particles that remained bound to the cell surface, and the cells were lysed by freeze-thawing. Viral lysates were centrifuged at 105,000 × <u>g</u> for 1h at 4°C in a 45Ti rotor using a Beckman ultracentrifuge (OptimaL-90). The supernatant was discarded and the viral particles in the pellet were resuspended in 4 ml TNC buffer (10 mM Tris, pH7.5, 140 mM NaCl, 10 mM CaCl_2_), extracted with trichloromonofluoromethane (Genetron, CYDSA 11, Mexico City, Mexico) and placed on top of a 1 ml 40% sucrose cushion in TNC. The samples were centrifuged at 195,000 × <u>g</u> for 2 h at 4°C in a SW55Ti rotor. Finally, the resulting pellets containing the semi-purified viral particles were dissolved in TNC.

### Isopycnic gradient

MA104 cells were infected with RRV at an MOI of 5 in the presence of either 0.5 µg/ml BFA or 10µM GCA, and at 12 hpi the cells were lysed by freeze-thawing. The cell lysate was extracted with trichloromonofluoromethane as described above, and the aqueous phase was mixed with 2.2 g of CsCl and TNC buffer up to 5 ml. The samples were centrifuged at 158,000 × <u>g</u> for 18h at 4°C on a SW55Ti rotor (Beckman). Finally, the opalescent bands corresponding to TLPs and DLPs were collected by puncture with a syringe and stored at 4°C. Before use, the viral particles were desalted in a Sephadex G25 spin column.

### Transmission electron microscopy

MA104 cells were seeded onto sapphire discs and infected with RRV an MOI of 250 viroplasm forming units (VFU)/ml in the presence of 0.5 µg/ml BFA or 10µM GCA. At 6 hpi the cells were fixed with 2.5% glutaraldehyde in 100 mM Na/K-phosphate buffer, pH 7.4 for 1 h at 4°C and kept in 100 mM Na/K-phosphate buffer overnight at 4°C. Afterwards, samples were post-fixed in 1% osmium tetroxide in 100 mM Na/K-phosphate buffer for 1 h at 4°C, dehydrated in a graded ethanol series starting at 70% followed by two changes in acetone and embedded in Epon. Ultrathin sections (60-80 nm) were cut and stained with uranyl acetate and lead citrate. Images were acquired using a transmission electron microscope (CM12, Philips, Eindhoven, The Netherlands) equipped with a CCD camera (Ultrascan 1000, Gatan, Pleasanton, CA, USA) at an acceleration of 100kV and processed using ImageJ version 2.0.0-rc-69/1.52i (Creative Common, http://imageJ.net/contributors).

### Metabolic labeling of proteins

MA104 cells were pre-treated with BFA (0.5 µg/ml) or GCA (10 µM) for 30 min. Afterwards, the cells were mock-infected or infected with RRV (MOI of 5) in the presence of inhibitors for 1h. At the end of the adsorption period, the cells were washed, and fresh media containing the drugs were added. At 7 hpi, the cells were starved for 30 min in the methionine-free medium before pulse-labeling with 25 µCi/ml of Easy Tag Express ^35^S labeling mix (Perkin Elmer, Shelton, CT, USA) for 30 min. At 8 hpi, cells were washed three times, and the monolayers were incubated in complete medium before lysis in Laemmli sample buffer at 10 hpi. The radiolabeled proteins were resolved in a 10% SDS-PAGE, followed by autoradiography.

For ^3^H-mannose radiolabeling, cells were treated and infected as described above. At 6 hpi the cells were starved in a glucose-free medium for 30 min followed by labeling with 200 µCi/ml of mannose-D-[2-^3^H(N)] (Perkin Elmer, Shelton, CT) for 1.5 h. At 8 hpi, the ^3^H-labelling medium was removed, and the cells were incubated in complete medium before lysis in Laemmli sample buffer at 10 hpi. The radiolabeled proteins were resolved by 10% SDS-PAGE, followed by fluorography.

### Western blot analysis

Proteins in cell lysates were separated in a 10% SDS-PAGE and transferred to Immobilon NC (Millipore Merck, Darmstadt, Germany) membranes. The membranes were blocked with 5% nonfat dry milk in PBS for 1 h at room temperature and then incubated with primary antibodies diluted in PBS containing 0.1% nonfat dry milk. The membranes were then incubated with the corresponding secondary antibody conjugated to horseradish peroxidase. The peroxidase activity was developed using the Western Lightning Chemiluminescence Reagent Plus (PerkinElmer) according to the manufacturer’s instructions. The blots were also probed with an anti-vimentin antibody, which was used as a loading control.

### Immunofluorescence

MA104 cells grown on glass coverslips were pre-treated with BFA (0.5 µg/ml) or GCA (10 µM) for 30 min and then infected with rotavirus at an MOI of 1. Six hpi, cells were fixed with 2% paraformaldehyde in PBS for 20 min at room temperature. After this time, the cells were washed four times with 50 mM NH_4_Cl in PBS and permeabilized by incubation with 0.5% Triton X-100 for 15 min at room temperature. The coverslips were incubated in blocking buffer (1% bovine serum albumin) for 1h at room temperature (RT) and then with primary antibodies diluted in blocking buffer overnight at 4°C. The cells were then washed five times and incubated with the Alexa-labeled secondary antibodies in blocking buffer for 1 h at RT. Finally, cell nuclei were stained with 30 nM DAPI (4′,6-diamidino-2-phenylindole, Invitrogen, Eugene, OR, USA) for 5 min and the coverslips were mounted onto glass slides with CitiFluor AF1 antifading (Electron Microscopy Sciences, Emsdiasum, Hatfield Penn). Images were acquired with an inverted 3I spinning disk confocal microscope (Zeiss Observer Z.1) coupled to a digital EMCCD Andor Ixon (512×512 pixels) and processed using Fiji ImageJ 1.52n(89).

### Statistical analysis

Statistical significance was evaluated using a one-way analysis of variance (ANOVA) test followed by Tukey’s multiple comparison posttest or an unpaired t-test using GraphPad Prism 6.01 (GraphPad Software Inc.).

## ACKNOWLEDGMENTS

J.L.M. is recipient of a scholarship from CONACyT. This work was partially supported by grant IG200317 from DGAPA/UNAM, by grant A1-5-15356 from the National Council for Science and Technology, Conacyt-Mexico, and by the University of Zurich. The funders had no role in study design, data collection, and interpretation, or the decision to submit the work for publication.

---

**Fig. S1.**
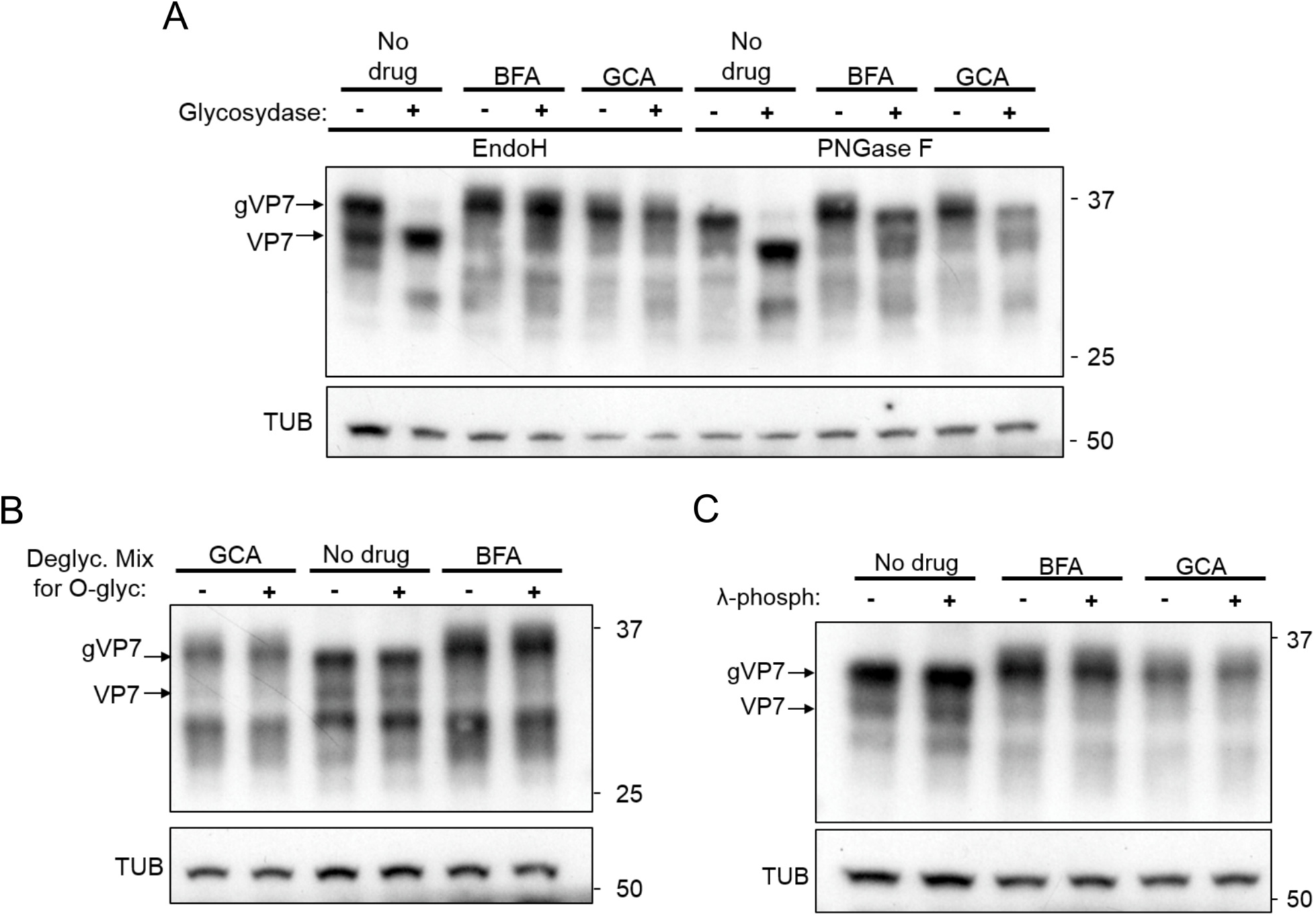
BFA- and GCA-induced modification of VP7 electrophoretic mobility is related neither to N-/O-glycosylation nor to phosphorylation. To investigate the modification of the electrophoretic migration of the RRV VP7 protein induced by BFA and GCA, a recombinant VP7 protein was overexpressed in MA104 cells; BFA or GCA (5 μg/ml) were added at 1 hpt and cells were lysed (7.6% SDS, 125 mM TrisHCl, pH 6.8) at 18 hpt. Cell extracts underwent treatment with either of the following enzymes according to the manufacturer instructions (Life Technologies): PNGaseF (Peptide-*N*-Glycosidase F), which removes almost all types of *N*-linked (Asn-linked) glycosylation; EndoH (endo-β-N-acetylglucosaminidase-H), which removes only high mannose and some hybrid types of *N*-linked carbohydrates; a Protein Deglycosylation Mix, which in addition to all N-linked glycans removes many common O-linked glycans; λ-phospatase, a protein phosphatase with activity towards phosphorylated serine, threonine and tyrosine residues. The figure shows a representative Western blot of the cell extracts incubated with the enzymes indicated. The glycosylated (gVP7) and deglycosylated (VP7) forms of VP7 are indicated. Following cell treatment with BFA/GCA, VP7 becomes EndoH-resistant and part of VP7 also becomes PNGaseF-resistant (A), suggesting that the modification induced by treatment with BFA/GCA is not N-glycosylation. Also, it is neither O-glycosylation nor phosphorylation, as shown in panels (B) and (C), respectively. The bands revealed by anti-VP7 antibody at a lower MW than that of VP7 (particularly intense with the Protein Deglycosylation Mix) most likely represent VP7 fragments derived from degradation occurred during incubation of the cell extracts with the enzymes. Tubulin (TUB) was used as loading control.

**Fig. S2.**
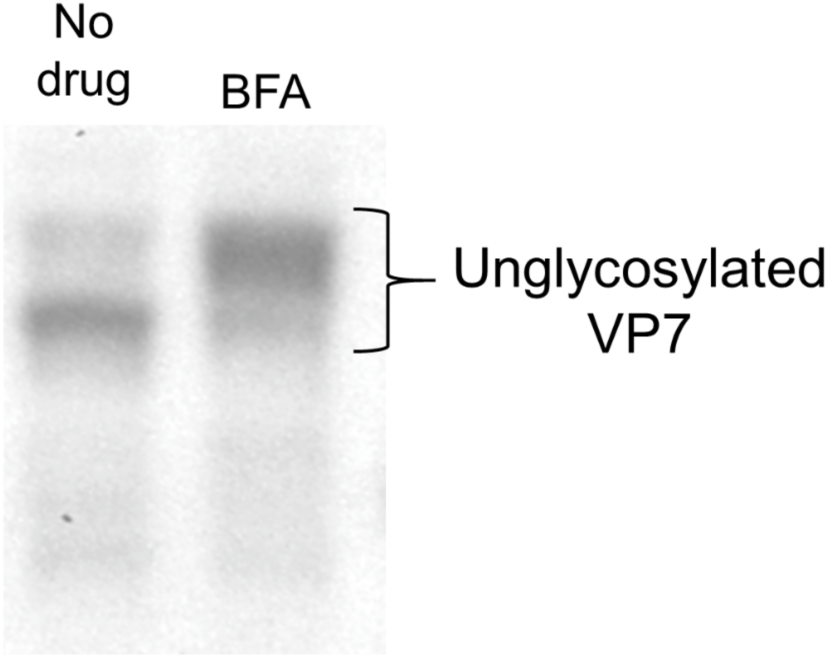
BFA modified the electrophoretic mobility of an unglycosylated form VP7. A recombinant RRV VP7 protein with the N-glycosylation site mutated (69-NST-71 to 69-QSG-71) was overexpressed in MA104 cells; BFA (2.5 μg/ml) were added at 1 hpt and the cells were lysed (7.6% SDS, 125 mM TrisHCl pH 6.8) at 18 hpt. The figure shows a representative Western blot analysis of cells transfected with the unglycosylated VP7. The expression of VP7 was detected with a specific antibody.

